# Embedding to Reference t-SNE Space Addresses Batch Effects in Single-Cell Classification

**DOI:** 10.1101/671404

**Authors:** Pavlin G. Poličar, Martin Stražar, Blaž Zupan

## Abstract

Dimensionality reduction techniques, such as t-SNE, can construct informative visualizations of high-dimensional data. When working with multiple data sets, a straightforward application of these methods often fails; instead of revealing underlying classes, the resulting visualizations expose data set-specific clusters. To circumvent these batch effects, we propose an embedding procedure that takes a t-SNE visualization constructed on a reference data set and uses it as a scaffold for embedding new data. The new, secondary data is embedded one data-point at the time. This prevents any interactions between instances in the secondary data and implicitly mitigates batch effects. We demonstrate the utility of this approach with an analysis of six recently published single-cell gene expression data sets containing up to tens of thousands of cells and thousands of genes. In these data sets, the batch effects are particularly strong as the data comes from different institutions and was obtained using different experimental protocols. The visualizations constructed by our proposed approach are cleared of batch effects, and the cells from secondary data sets correctly co-cluster with cells from the primary data sharing the same cell type.

## 1 Introduction

Two-dimensional embeddings and their visualizations may assist in the analysis and interpretation of high-dimensional data. Intuitively, two data instances should be co-located in the resulting visualization if their multi-dimensional profiles are similar. For this task, non-linear embedding techniques such as t-distributed stochastic neighbor embedding (t-SNE) [1] or uniform manifold approximation and projection [2] have recently complemented traditional data transformation and embedding approaches such as principal component analysis (PCA) and multi-dimensional scaling [3, 4]. While useful for visualizing data from a single coherent source, these methods may encounter problems if the data comes from multiple sources. Here, when performing dimensionality reduction on a merged data set, the resulting visualizations would typically reveal source-specific clusters instead of grouping data instances of the same class-type, regardless of data sources. This source-specific confounding is often referred to as *domain shift* [5], *covariate shift* [6] or *dataset shift* [7]. In bioinformatics, the domain-specific differences are more commonly referred to as *batch effects* [8–10].

Massive, multi-variate biological data sets often suffer from these source-specific biases. Consider an example from single-cell genomics, the domain we will focus on in this manuscript and that was — besides current scientific challenges — selected also due to the availability and abundance of recently published data. Single-cell RNA sequencing (scRNA-seq) data sets are the result of isolating RNA molecules from individual cells, which serve as an estimate of the expression of cell’s genes. Single-cell studies, which can exceed thousands of cells and tens of thousands of genes, typically start with the analysis of cell types. Here, it is generally expected that cells of the same type would cluster together in two-dimensional data visualisation [10]. For instance, Fig. 1.a shows t-SNE embedded data from mouse brain cells originating from the visual cortex [11] and the hypothalamus [12]. The figure reveals distinct clusters but also separates the data from the two brain regions. These two regions share the same cell types and — contrary to the depiction in Fig. 1.a — we would expect the data points from the two studies to overlap. Batch effects similarly prohibit the utility of t-SNE in the exploration of pancreatic cells in Fig. 1.b, which renders the data from a pancreatic cell atlas [13] and similarly-typed cells from diabetic patients [14]. Just like with data from brain cells, pancreatic cells cluster primarily by data source, again resulting in an uninformative visualization driven by batch effect.

**Fig. 1.**
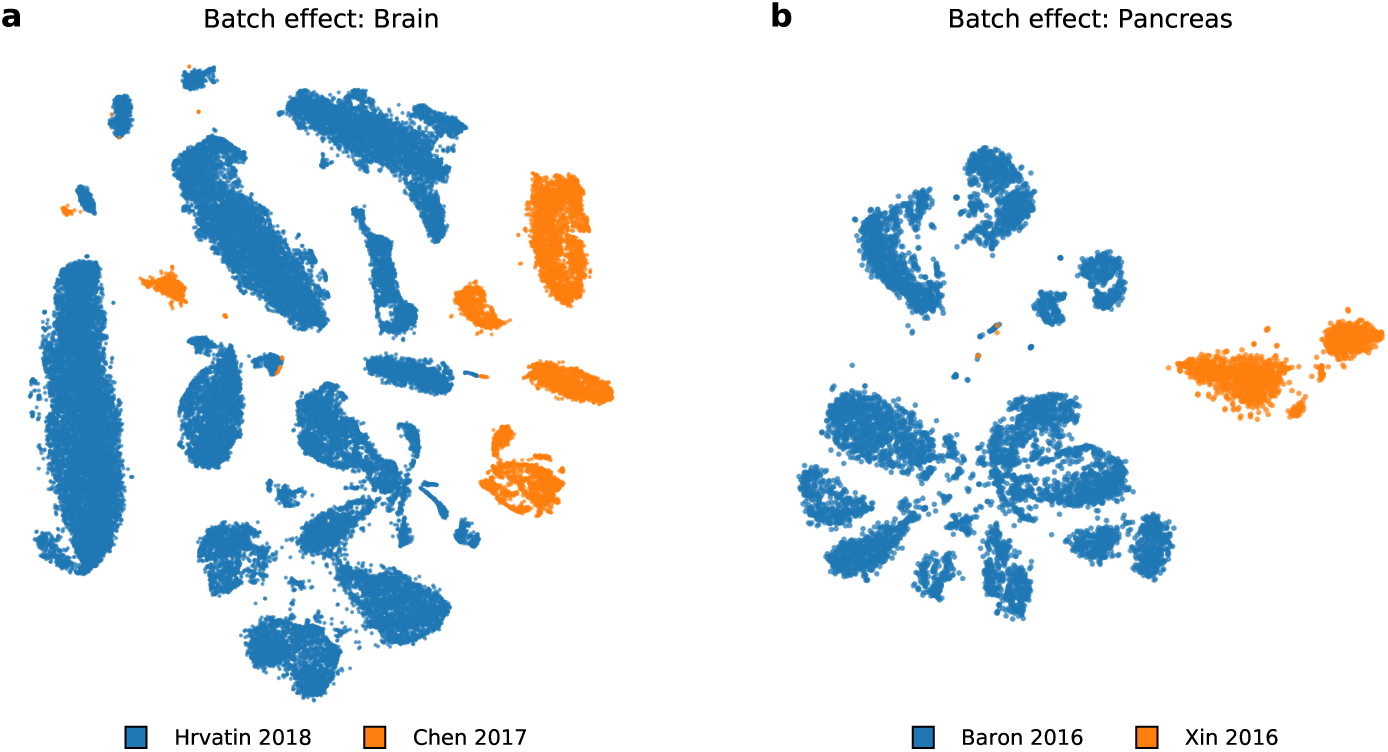
Batch effects are a driving factor of variation between the data sets. Depicted is a t-SNE visualisation of two pairs of data sets. In each pair, the data sets share cell types, so it would be expected that the cells from the reference data (blue) would mix with the cells in a secondary data sets (orange). Instead, t-SNE visualisation clusters data according to the data source.

Current solutions to embedding the data from various data sources address the batch effect problems up-front. The data is typically preprocessed and transformed such that the batch effects are explicitly removed. Recently proposed procedures for batch effect removal include canonical correlation analysis [8] and mutual nearest-neighbors [9, 10]. In these works, batch effects are deemed removed when cells from different sources exhibit good mixing in a t-SNE visualization. The elimination of batch effects may require aggressive data preprocessing which may blur the boundaries between cell types. Another problem is also the inclusion of any new data, for which the entire data analysis pipeline must be rerun, usually resulting in a different layout and clusters that have little resemblance to original visualization and thus require reinterpretation.

We propose a direct solution of rendering t-SNE visualizations that addresses batch effects. Our approach treats one of the data sets as a *reference* and aims to embed the cells from another, *secondary data set* to a common low-dimensional space. We construct a t-SNE embedding using the reference data set, and then use it as a scaffold for the embedding of data points from the secondary data. The key idea underpinning our approach is that the embedding is performed one data point at a time. Independence of each new embedded data instance from the secondary data set causes the clustering landscape to depend only on the reference scaffold, thus removing data source-driven variation. In other words, when including new data, the scaffold inferred from the reference data set is kept unchanged and defines a “gravitational field”, independently driving the embedding of each new instance. For example, in Fig. 2, the cells from the visual cortex define the scaffold (Fig. 2.a) into which we embed the cells from the hypothalamus (Fig. 2.b). Unlike in their joint t-SNE visualization (Fig. 1.a), the hypothalamic cells are dispersed across the entire embedding space and their cell type correctly matches the prevailing type in reference clusters.

**Fig. 2.**
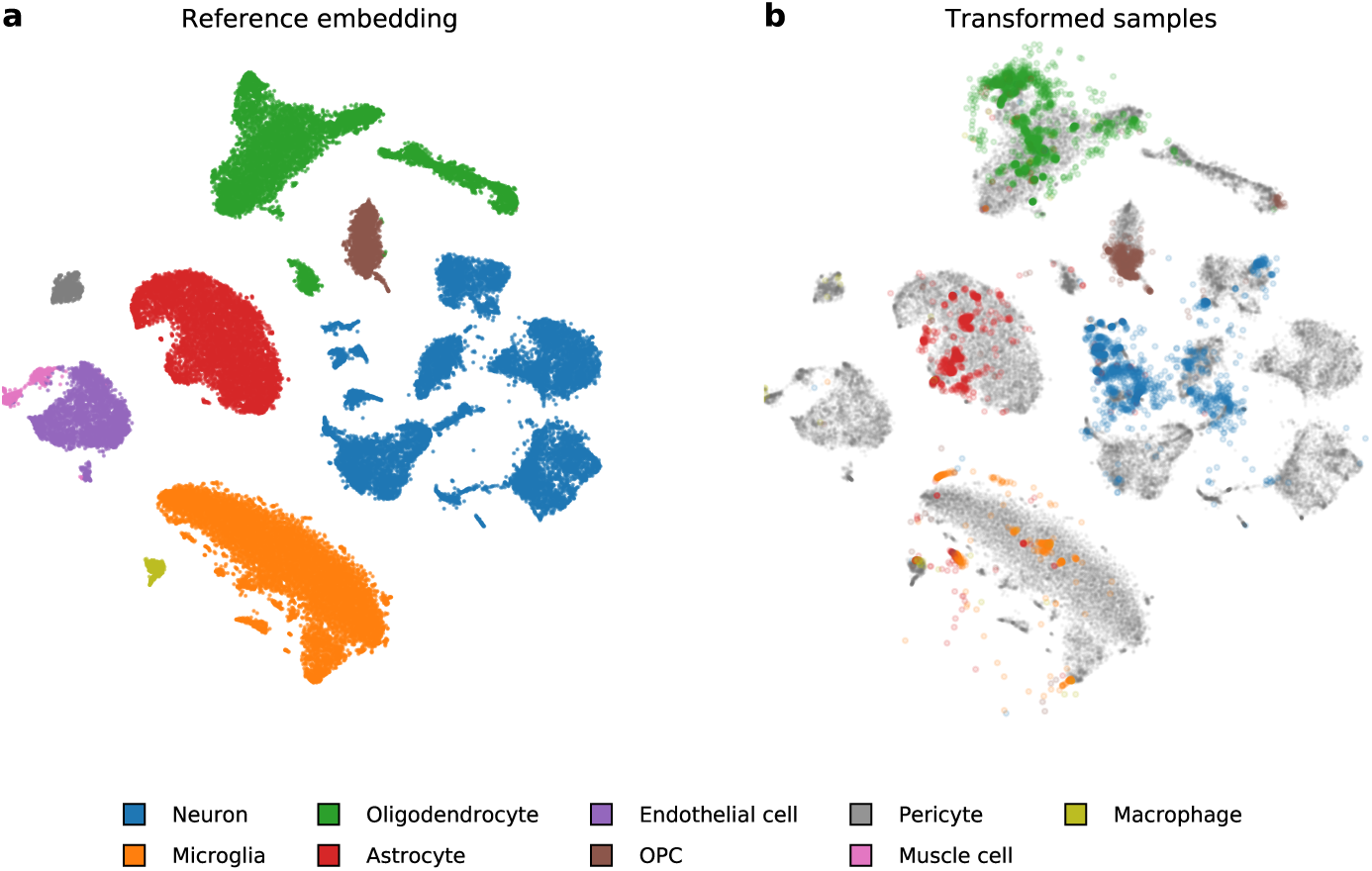
A two-dimensional embedding of a reference containing brain cells (a) and the corresponding mapping of secondary data containing hypothalamic cells (b). Notice that the majority of hypothalamic cells were mapped to their corresponding reference cluster. For instance, astrocyte cells marked with red on the right were mapped to an oval cluster of same-typed cells denoted with the same color in the visualization on the left.

The proposed solution is implemented using a mapping of new data to an existing t-SNE visualization. While the utility of such an algorithm was already hinted at in recent publication [15], here we provide its practical and theoretically-grounded implementation. Considering the abundance of recent publications on batch effect removal, we present surprising evidence that a computationally more direct and principled embedding procedure solves the batch effects problem when constructing interpretable visualizations from different data sources.

## 2 Methods

We describe an end-to-end pipeline that uses fixed t-SNE coordinates as a scaffold for embedding new (secondary) data, enabling joint visualisation of multiple data sources while mitigating batch effects. Our proposed approach starts by using t-SNE to embed a reference data set, with the aim of constructing a two-dimensional visualisation to facilitate interpretation and cluster classification. Then, the placement of each new sample is optimized independently via the t-SNE loss function. Independent treatment of each data instance from a secondary data set disregards any interactions present in that data set, and prevents the formation of clusters that would be specific to the secondary data. Below, we start with a summary of t-SNE and its extensions (Sec. 2.1, introducing the relevant notation, upon which we base our secondary data embedding approach (Sec. 2.2).

### 2.1 Data embedding by t-SNE and its extensions

t-SNE is a local, non-linear dimensionality reduction method, tailored to the visualisation of high-dimensional data sets. Given a multi-dimensional data set **X** = {**x**_1_, **x**_2_, …, **x**_*N*_} ∈ ℝ^*D*^ where *N* is the number of samples in the reference data set, t-SNE aims to find a low dimensional embedding **Y** = {**y**_1_, **y**_2_, …, **y**_*N*_} ∈ ℝ^*d*^ where *d* « *D*, such that if points **x**_*i*_ and **x**_*j*_ are close in the multi-dimensional space, their corresponding embeddings **y**_*i*_ and **y**_*j*_ are also close. Since t-SNE is primarily used as a visualization tool, *d* is typically set to two. The similarity between two data points in t-SNE is defined as:

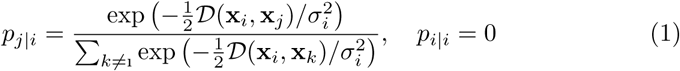

where *𝒟* is a distance measure. This is then symmetrized to

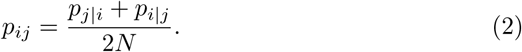

The bandwidth of each Gaussian kernel *σ*_*i*_ is selected such that the perplexity of the distribution matches a user-specified parameter value

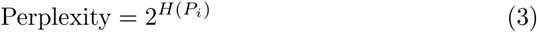

where *H*(*P*_*i*_) is the Shannon entropy of *P*_*i*_,

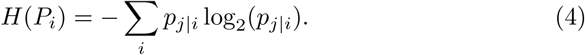

Different bandwidths *σ*_*i*_ enable t-SNE to adapt to the varying density of the data in the multi-dimensional space.

The similarity between points **y**_*i*_ and **y**_*j*_ in the embedding space is defined using the *t*-distribution with one degree of freedom

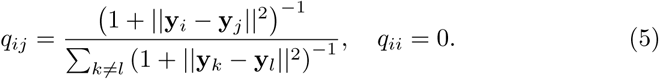

The t-SNE method finds an embedding **Y** that minimizes the Kullback-Leibler (KL) divergence between **P** and **Q**,

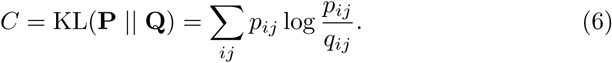

The time complexity needed to evaluate the similarities in Eq. 5 is 𝒪(*N*^2^), making the application of the algorithm impractical for any reasonably-sized data. To address larger data sets, we adopt a recent approach for low-rank approximation of gradients based on polynomial interpolation which reduces the time complexity of t-SNE to 𝒪(*N*). This approximation enables the visualization of massive data sets, possibly containing millions of data points [16].

The resulting embeddings substantially depend on the value of the perplexity parameter. Perplexity can be interpreted as the number of neighbors for which the distances in the embedding space are preserved. Small values of perplexity result in tightly-packed clusters of points and effectively ignore the long-range interactions between clusters. Larger values may result in a more globally consistent visualisations, preserving distances on a large scale and organizing clusters in a more meaningful way. Larger values of perplexity can lead to merging of multiple small clusters, thus obscuring local aspects of the data [15].

The trade-off between the local organization and global consistency may be achieved by replacing the Gaussian kernels in Eq. 1 with a mixture of Gaussians of varying bandwidths [17]. Multi-scale kernels are defined as

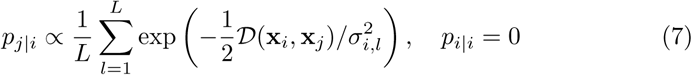

where *L* is the number of mixture components. The bandwidths *σ*_*i,l*_ are selected in the same manner as in Eq. 1, but with a different value of perplexity for each *l*. In our experiments, we used a mixture of two Gaussian kernels with perplexity values of 50 and 500. We note that a similar formulation of multi-scale kernels was proposed in [15], and we found the resulting embeddings are visually very similar to those obtained with the approach described above (not shown for brevity).

### 2.2 Adding new data points to reference embedding

Our algorithm, which embeds new data points to a reference embedding, consists of estimating similarities between each new point and the reference data and optimizing the position of each new data point in the embedding space. Unlike parametric models such as principal component analysis or autoencoders, t-SNE does not define an explicit mapping to the embedding space, and embeddings need to be found through loss function optimization.

The position of a new data point in embedding space is initialized to the median reference embedding position of its *k* nearest neighbors. While we found the algorithm to be robust to choices of *k*, we use *k* = 10 in our experiments.

We adapt the standard t-SNE formulation from Eqs. 1 and 5 with

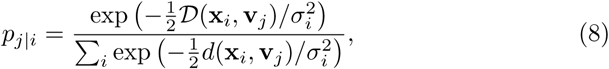

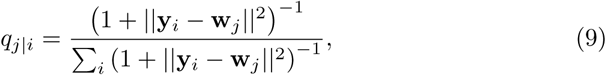

where **V** = {**v**_1_, **v**_2_, …, **v**_*M*_} ∈ ℝ^*D*^ where *M* is the number of samples in the new data set and **W** = {**w**_1_, **w**_2_, …, **w**_*M*_} ∈ ℝ^*d*^. Additionally, we omit the symmetrization step in Eq. 2. This enables new points to be inserted into the embedding independently of one another. The gradients of **w**_*j*_ with respect to the loss (Eq. 6) are:

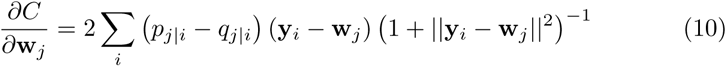

In the optimization step, we refine point positions using batch gradient descent. We use an adaptive learning rate scheme with momentum to speed up the convergence, as proposed by Jacobs [18, 19]. We run gradient descent with momentum *α* set to 0.8 for 250 iterations, where the optimization converged in all our experiments. The time complexity needed to evaluate the gradients in Eq. 10 is *𝒪* (*N · M*), however, by adapting the same polynomial interpolation based approximation, this is reduced to *𝒪* (*N*).

Special care must be taken to reduce the learning rate *η* as the default value in most implementations (*η* = 200) may cause points to “shoot off” from the reference embedding. This phenomenon is caused due to the embedding to a previously defined t-SNE space, where the distances between data points and corresponding gradients of the optimization function may be quite large. When running standard t-SNE, points are initialized and scaled to have variance 0.0001. The resulting gradients tend to be very small during the initial phase, resulting in stable convergence. When embedding new samples, the span of the embedding is much larger, resulting in substantially larger gradients, and the default learning rate causes points to move very far from the reference embedding. In our experiments, we found that decreasing the learning rate to *η* ∼ 0.1 produces stable solutions. This is especially important when using the interpolation-based approximation, which places a grid of interpolation points over the embedding space, where the number of grid points is determined by the span of the embedding. Clearly, if even one point “shoots off” far from the embedding, the number of required grid points may grow dramatically, increasing the runtime substantially. The reduced learning rate suppresses this issue, and does not slow the convergence because of the adaptive learning rate scheme, provided the optimization is run for a sufficient number of steps.

## 3 Experiments and Discussion

We apply the proposed approach to t-SNE visualizations of single-cell data. Data in this realm include a variety of cells from specific tissues, and characterize them through gene expression. In our experiments, we considered several recently published data sets where cells were annotated with the cell type. Our aim was to construct t-SNE visualizations where similarly-typed cells would cluster together, despite systematic differences between data sources. Below, we list the data sets, describe single-cell specific data preprocessing procedures, and display the resulting data visualizations. Finally, we discuss the success of the proposed approach in alleviating the batch effects.

### 3.1 Data

We use three pairs of reference and secondary single-cell data sets originating from different organisms and tissues. The data in each pair were chosen so that the majority of cell types from the secondary data set were included in the reference set (Table 1).

**Table 1.**
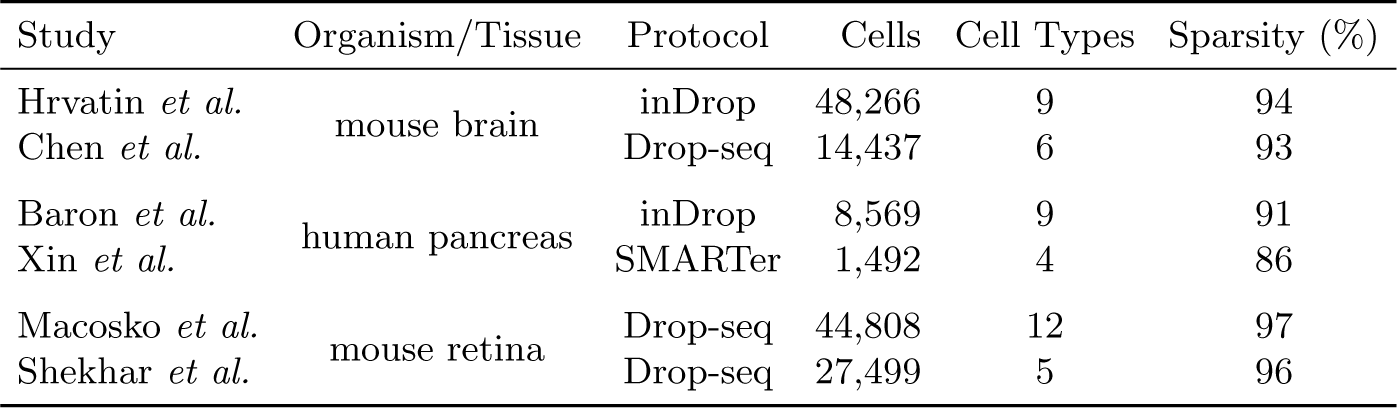
Data sets used in our experiments. In each pair, the first data set (Hrvatin *et al*., Baron *et al*., and Macoscko *et al*.) was used as a reference. In all cases, we relied on the quality control and annotations from the original publication. To facilitate comparisons, the cell annotations were harmonized using cell type annotations from the cell ontology [20]. Notice that different RNA sequencing protocols were used to estimate gene expressions. We report the number of cell types from each data set retained after preprocessing. Single-cell data is sparse, typically containing less than 10% expressed genes per cell.

| Study | Organism/Tissue | Protocol | Cells | Cell Types | Sparsity (%) |
| --- | --- | --- | --- | --- | --- |
| Hrvatin *et al.* | mouse brain | inDrop | 48,266 | 9 | 94 |
| Chen *et al.* |  | Drop-seq | 14,437 | 6 | 93 |
| Baron *et al.* | human pancreas | inDrop | 8,569 | 9 | 91 |
| Xin *et al.* |  | SMARTer | 1,492 | 4 | 86 |
| Macosko *et al.* | mouse retina | Drop-seq | 44,808 | 12 | 97 |
| Shekhar *et al.* |  | Drop-seq | 27,499 | 5 | 96 |

The cells in the data sets originates from the following three tissues:

#### Mouse brain

The data set from Hrvatin *et al*. [11] contains cells from the visual cortex exploring transcriptional changes after exposure to light. This was used as a reference for the data from Chen *et al*. [12], containing various cells from the mouse hypothalamus and their reaction to food deprivation. From the secondary data, we removed cells with no corresponding types in the reference, namely ependymal cells, epithelial cells, tanycytes, and unlabelled cells.

#### Human pancreas

The data set from Baron *et al*. [13] was created as an atlas of pancreatic cell types. We used this set as a reference for data from Xin *et al*. [14], who examined transcriptional differences between healthy and type 2 diabetic patients.

#### Mouse retina

The data set from Macosko *et al*. [21] was created as an atlas of mouse retinal cell types. We used this as a reference for the data from Shekhar *et al*. [22], who built an atlas for different types of retinal bipolar cells.

### 3.2 Single-cell data preprocessing pipeline

Due to the specific nature of single-cell data, additional steps must be taken to properly apply t-SNE. We use a standard single-cell preprocessing pipeline, consisting of the selection of 3,000 representative genes (see Sec. 3.3), library size normalization, log-transformation, standardization, and PCA-based representation that retains 50 principal components [10, 23]. To obtain the reference embedding, we apply multi-scale t-SNE using PCA initialization [15]. Due to high-dimensionality of the preprocessed input data we use cosine distance to estimate similarities between reference data points [24]. When adding new data points from the secondary data set to the reference embedding, we select 1,000 genes present in both data sets and use these to estimate the similarities between the secondary data item and reference data points. The similarities are estimated using cosine similarity. We note that similarities are computed using the raw count matrices. The preprocessing stages are detailed in accompanying Python notebooks (Sec. 3.5).

### 3.3 Gene selection

Single-cell data sets suffer from high levels of technical noise and low capture efficiency, resulting in sparse expression matrices [25]. To address this problem, we use a specialized feature-selection method, which exploits the mean-dropout relationship of expression counts as recently proposed by Kobak and Berens [15]. Here, genes with higher than expected dropout rate are regarded as potential markers for cell subpopulations and are retained in the data.

Given an expression matrix **X** ∈ ℝ^*N* ×*G*^ where *N* is the number of samples and *G* is the number of genes in the data set, we compute the fraction of cells where a gene *g* was not expressed

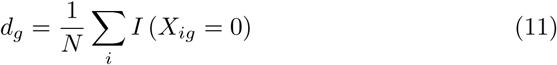

The mean log_2_ expression of the genes is computed from all the cells where gene was expressed:

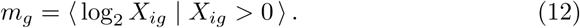

All genes expressed in less than ten cells are discarded. In order to select a specific number of *Ĝ* genes, we use a binary search to find a value *b* such that

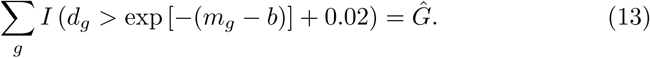

### 3.4 Results and Discussion

Figs. 2, 3, and 4 show the embeddings of the reference data sets and their corresponding embeddings of the secondary data sets. In all the figures, the cells from the secondary data sets were positioned in the cluster of same-typed reference cells, providing strong evidence of the success of the proposed approach. There are some deviations to these observations; for instance, in Fig. 2 several oligodendrocyte precursor cells (OPCs) were mapped to oligodendrocytes. This may be due to differences in annotation criteria by different authors, or due to inherent similarities of these types of cells. Examples of such erroneous placements can be found in other figures as well, but they are not common and constitute less then 5% of the cells (less than 5% in brain, %1 in pancreas and %2 in retina secondary data sets).

**Fig. 3.**
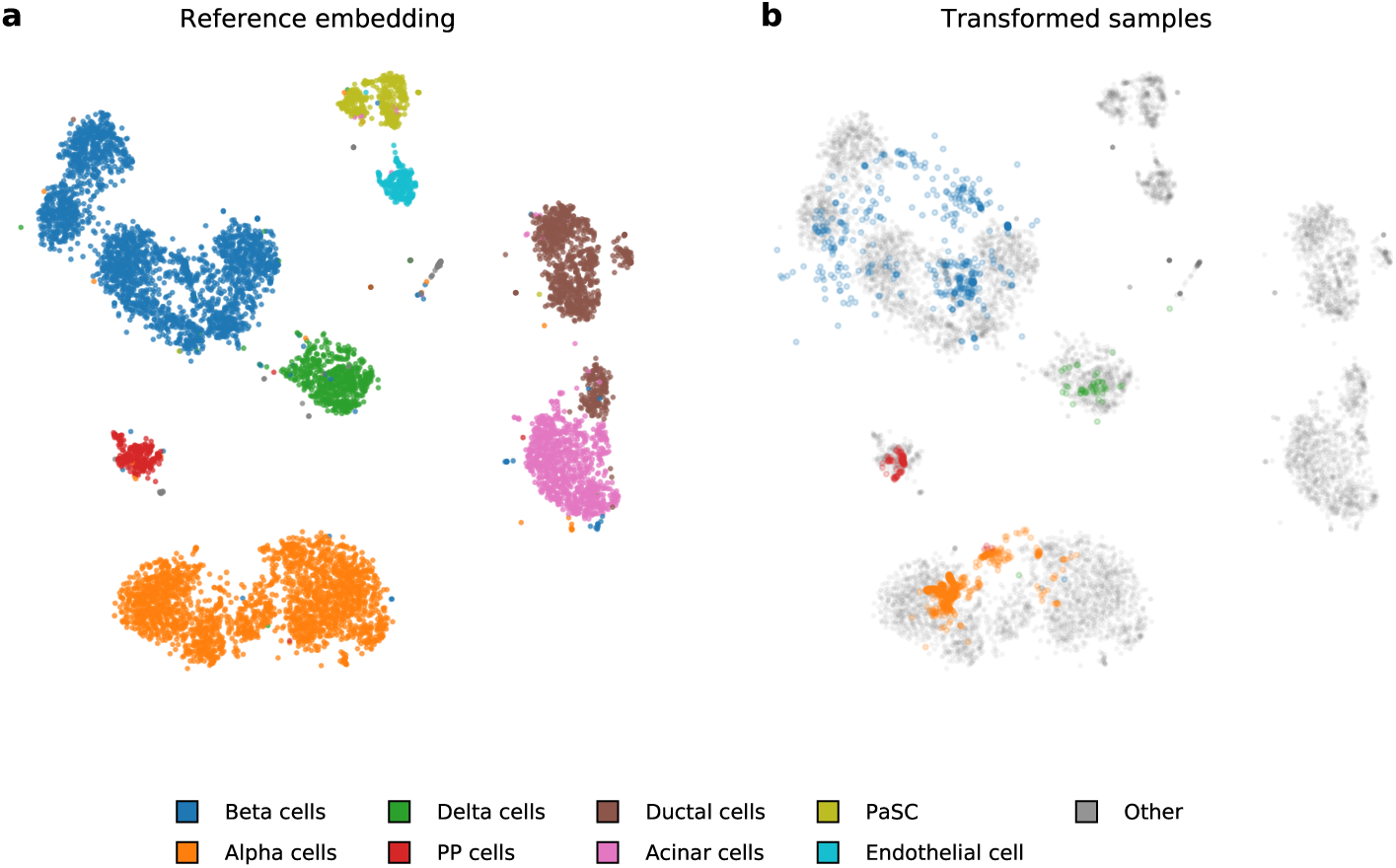
Embedding of pancreatic cells from Baron *et al*. [13] and cells from the same tissue from Xin *et al*.. [14]. Just like in Fig. 2 the vast majority of the cells from the secondary data set were correctly mapped to the same-typed cluster of reference cells.

**Fig. 4.**
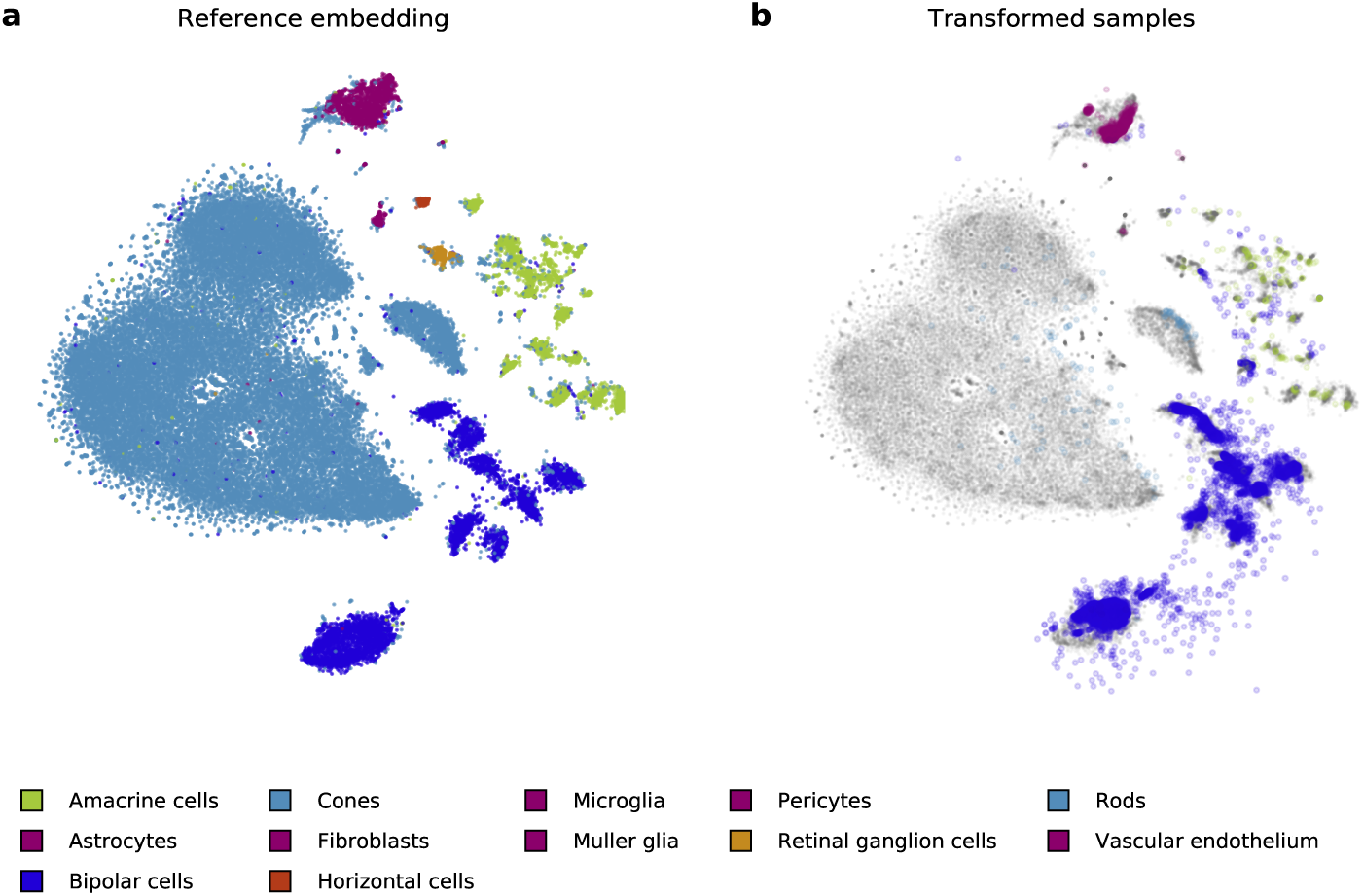
An embedding of a large reference of retinal cells from Macosco *et al*. [21] (a) and mapping of cells from a smaller study that focuses on bipolar cells from Shekhar *et al*. [22] (b). We use colors consistent with the study by Macosko *et al*..

Notice that we could simulate the split between reference and secondary data sets using one data set only and perform cross-validation, however this type of experiment would not incorporate batch effects. We want to remind the reader that handling batch effects were central to our endeavour and that the disregard of this effect could lead to overly-optimistic results and data visualizations strikingly different from ours. For example, compare the visualisations from Fig. 1.a and Fig. 2.b, or Figs. 1.b and 3.b.

There are additional modifications that we use in the embedding of the secondary data set that were recently proposed and enhance the original t-SNE visualization. One important extension is the use of multi-scale similarities that, besides local ordering of the data points, includes global optimization of cluster placement. For illustration, consider visualizations with standard and multi-scale t-SNE in Fig. 5. Notice, for instance, that in multi-scale t-SNE (Fig. 5.b) the clusters with neuronal cells are clumped together, while their placement in standard t-SNE is arbitrary (Fig. 5.a).

**Fig. 5.**
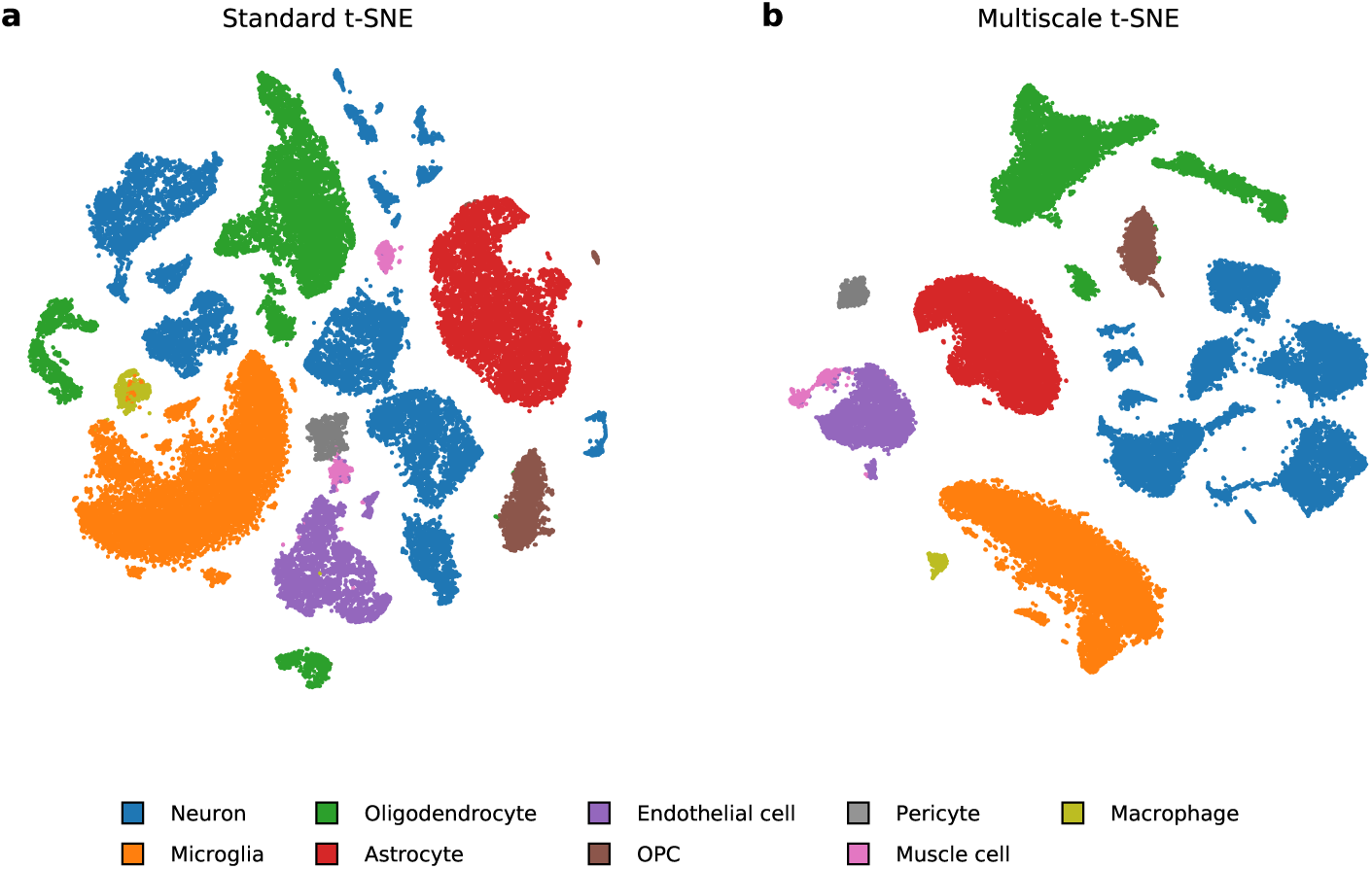
A comparison of standard and multi-scale t-SNE on data from the mouse visual cortex [11]. **(a)** Standard t-SNE places clusters arbitrarily. **(b)** Augmenting t-SNE with multi-scale similarities and using proper initialization provides a more meaningful layout of the clusters. Neuronal types occupy one region of the space. Oligodendrocyte precursor cells (OPCs) are mainly progenitors to oligodendrocytes, but may also differentiate into neurons or astrocytes.

We also observed the important role of gene selection in crafting the reference embedding spaces. We found that when selecting an insufficient number of genes, the resulting visualizations display overly-fragmented clusters. When the selection is too broad and includes lowly expressed genes, the subclusters tend to overlap. These effects can all be attributed to sparseness of the data sets and may be intrinsic to single-cell data. In our studies, we found that selection of 3,000 genes yields most informative visualizations (Fig. 6).

**Fig. 6.**
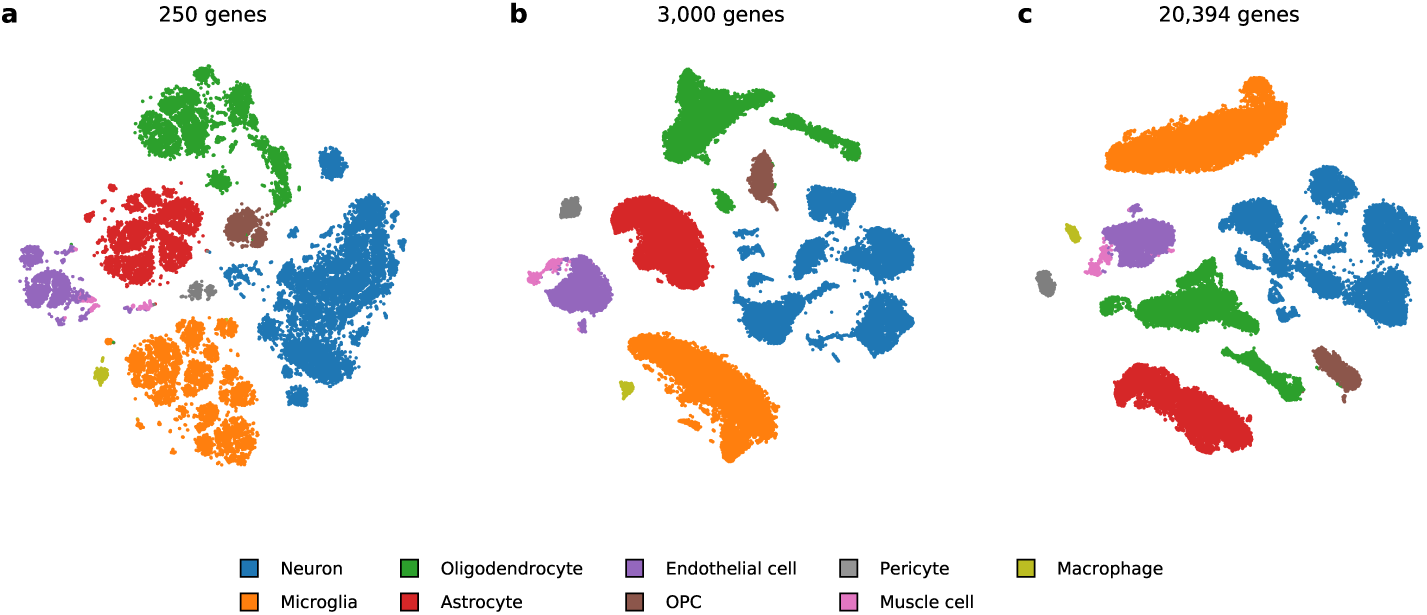
Gene selection plays an important role when constructing the reference embedding. **(a)** Using too few genes results in fragmented clusters. **(b)** Using an intermediate number of genes reveals clustering mostly consistent with cell annotations. **(c)** Including all the genes may lead to under-clustering of the more specialized cell types.

In principle, our theoretically-grounded embedding of secondary data into the scaffold defined by the reference embedding could be simplified with the application of the nearest neighbors-based procedure. For example, while describing a set of tricks for t-SNE, Kobak and Berens [15] proposed positioning new points into a known embedding by placing them in the median position of their 10 nearest neighbors, where the neighborhood was estimated in the original data space. Notice that we use this approach as well, but only for the initialization of positions of new data instances that are subject to further optimization. In Fig. 7 we demonstrate that nearest neighbor-based positioning is insufficient and may yield clumped visualizations where the optimal positioning using the t-SNE loss function is much more dispersed and rightfully shows a more considerable variation in the secondary data. Some data points may also fall into the empty regions between different clusters, while after optimization they typically move closer to same-typed groups.

**Fig. 7.**
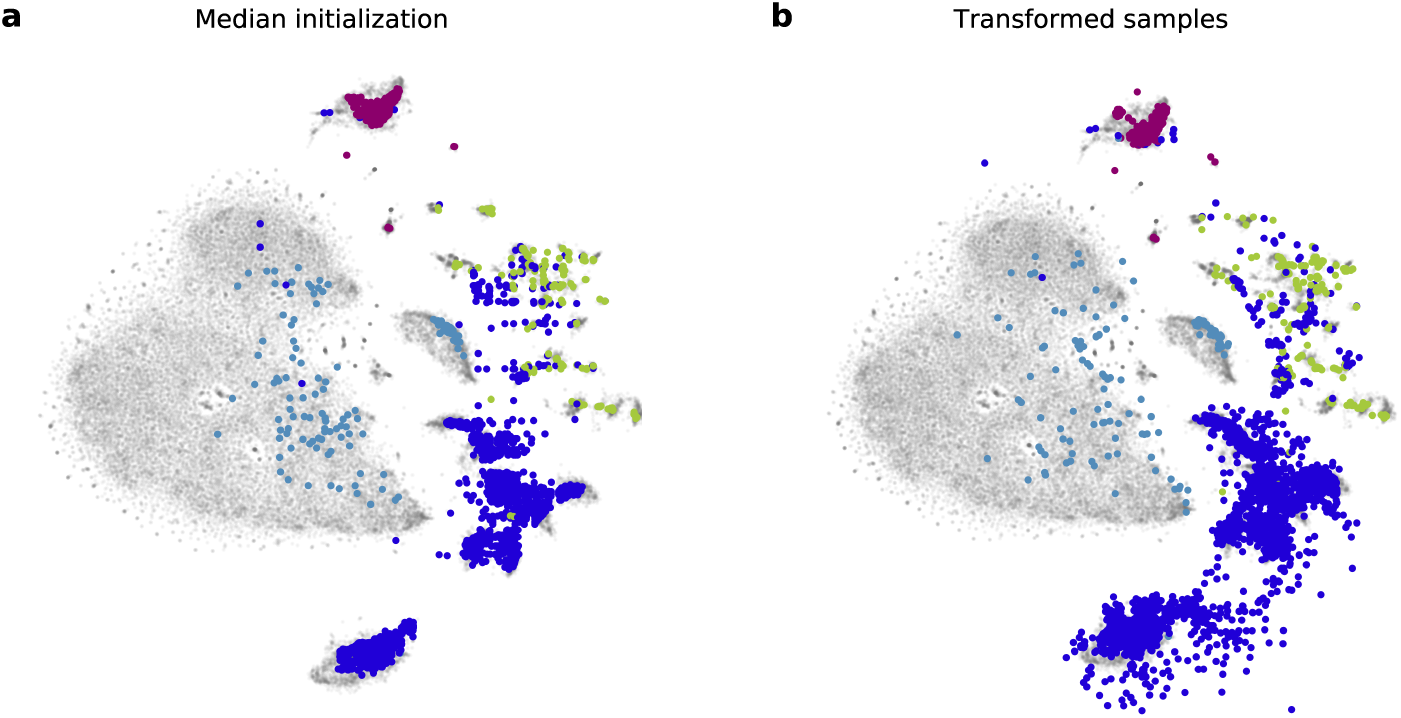
Comparison of data placement using the nearest neighbors approach from Kobak and Berens [15] and the optimized placement using our algorithm. **(a)** Data points are placed to the median position of their 10 nearest neighbors in the reference set. **(b)** Point positions are optimized, revealing a different, more dispersed placement that better reflects the variety of cells in the secondary data set.

The proposed method assumes that all cell types from the secondary data set are present in the reference. The proposed method would fail to reveal novel cell types in the secondary data set, possibly positioning them arbitrarily close to unrelated clusters. Procedures such as scmap were recently proposed to cope with such cases and identify the cells whose type is new and not included in the reference [26]. Our procedure does not address such cases, and for scaling-up to a wider collection of cell types relies on emerging availability of large collections of the reference data such as those managed by Human Cell Atlas initiative [27].

### 3.5 Implementation

The procedures described in this paper are provided as Python notebooks that are, together with the data, available in an open repository ^3^. All experiments were run using openTSNE ^4^, our open and extensible t-SNE library for Python.

## 4 Conclusion

Almost all recent publications of single-cell studies begin with a two-dimensional visualization of the data that reveals the cellular diversity containing many different cell-types from the study. While any dimensionality reduction technique can be used to render such a visualization, different variants of t-SNE are most often used. Due to the ability to explore biological mechanisms at the individual cell level, single-cell studies are increasingly widespread, and their publications in the past couple of years are abundant. One of the central tasks in single-cell studies is the classification of new cells based on findings from previous studies. Such transfer of knowledge is often difficult due to batch effects present in data from different sources. Addressing batch effects by adapting and extending t-SNE, the prevailing method used to present single-cell data in two-dimensional visualization motivated the research presented in this paper.

Our proposed approach uses a t-SNE embedding as a scaffold for the positioning of new cells within the visualization, and possibly for aiding in their classification. The three case studies incorporating pairs of data sets from different domains but with similar classifications demonstrate that our proposed procedure can effectively deal with batch effects to construct visualizations that correctly map secondary data sets onto a reference data set from an independent study that possibly uses different experimental protocol. While we focused here on reference visualizations constructed using t-SNE, this approach can be applied using any existing two-dimensional visualization.

## Acknowledgements

This work was supported by the Slovenian Research Agency Program Grant P2-0209, and by the BioPharm.SI project supported from European Regional Development Fund and the Slovenian Ministry of Education, Science and Sport. We would also like to thank Dmitry Kobak for discussions on t-SNE.

## Footnotes

3 https://github.com/biolab/tsne-embedding

^4^https://github.com/pavlin-policar/openTSNE

## Notes

https://github.com/biolab/tsne-embedding

